# Kinetics and efficacy of antibody drug conjugates in 3D tumour models

**DOI:** 10.1101/2023.02.14.528517

**Authors:** Chloe S Palma Chaundler, Haonan Lu, Ruisi Fu, Ning Wang, Hantao Lou, Gilberto Serrano de Almeida, Layla Mohammad Hadi, Eric O Aboagye, Sadaf Ghaem-Maghami

## Abstract

Antibody-drug conjugates (ADCs) are emerging targeted agents against cancer. Current studies of ADCs are performed on monolayer cultures which do not mimic the biophysical property of a tumour. Hence, *in vitro* models that can better predict the efficacy of ADCs *in vivo* are needed. In this study, we aim to optimise 3-dimentional cancer spheroid systems, which preserve the features of the tumour structure, to test the efficacy of two ADCs (T-DM1 and T-vcMMAE). Firstly, a set of reproducible spheroid models using epithelial ovarian cancer cell lines were established. Subsequently, phenotypic changes in spheroids were characterised upon ADC treatment. The penetration dynamics of ADCs into 3D tumour structure were also studied. Our data revealed that spheroids are less sensitive to ADCs compared to monolayer cultures. Interestingly, the small molecule component of ADCs-the cytotoxic payload-showed a similar decrease in efficacy in spheroids compared to monolayer cultures. Furthermore, we also gained new insight into ADC penetration dynamics and showed that ADCs can fully penetrate a tumour-like spheroid within 24h. The results suggest that although ADCs, as large molecule biological drugs, are likely to have slower penetration dynamics than small molecule compounds such as their cytotoxic payload, they could have comparable capability to kill cancer cells in 3D structures. This may be explained by the fact that multiple cytotoxic payloads are conjugated with each single antibody, which compensates the penetration deficiency of the large molecules. In conclusion, our work confirms that the tumour 3D structure could limit the therapeutic efficacy of ADCs. Nevertheless, optimising ADC design such as adjusting drug-to-antibody ratios could help to overcome this hurdle.

## INTRODUCTION

Ovarian cancer is a highly aggressive gynaecological disease that originates within the ovary or the fallopian tubes of women, ranking fifth in female cancer deaths around the globe (*1*). Whilst ovarian cancers comprise tumours of epithelial, germ and stem cell origin, epithelial ovarian cancers (EOC) are the most predominant subtype. In fact, EOC affects 90% of all ovarian cancer cases, leading to 140,000 annual deaths (*2*) (*3*). Treatment for advanced EOC patients still consists of the conventional therapeutic approaches used twenty years ago, involving debulking surgery and chemotherapy agents such as paclitaxel and carboplatin (*6*). Whilst these therapeutic approaches result in initial clinical remission rates of 75%, they are associated with 80% cancer recurrence (*7*). In fact, the median progression-free survival of the most common form of epithelial ovarian cancer, high grade serous ovarian cancer, does not surpass 16 to 21 months (*8*). Hence, there is a need to find more efficient treatments against EOC.

To overcome the challenges associated with current treatments against EOC, various targeted therapies which kill tumour cells more specifically, whilst limiting damage in normal cells are under development. Antibody drug conjugates (ADCs) are emerging classes of targeted agents which consist of a tumour antigen-specific monoclonal antibody attached to a range of cytotoxic payloads, such as DNA damaging or microtubule disrupting agents, through a chemical linker (*9*). ADCs have a rather clear mechanism of action: upon binding to tumour specific antigens, ADCs are internalised by cancer cells, followed by the release of their cytotoxic drugs mediated by low pH levels or the presence of specific proteases in the lysosome (*10*). Therefore, unlike conventional chemotherapy treatments, ADCs are able to exploit the distinct membrane proteome of cancer cells to deliver potent chemotherapeutic drugs specifically. Additionally, ADCs are also able to recruit and activate NK cells, further enhancing tumour killing through a mechanism known as antibody-dependent cellular cytotoxicity (ADCC) (*11*) (*12*). Given the advantages that ADCs pose over traditional chemotherapeutic approaches, there is an increasing interest in the development of ADCs for incurable cancers such as EOC. Currently, over 100 ADCs are in clinical development and eleven ADCs are approved by the US Food and Drug Administration (FDA) for use against solid tumours such as breast cancer (*13*).

Over the last decades, monoculture systems consisting of one or two cell types attached to plastic dishes have been the main *in vitro* models used to screen for novel anti-cancer drugs (*14*). The use of these simple models minimizes ethical issues and is more cost-effective and less time-consuming than *in vivo* models (*15*). Nevertheless, although several ADC agents against EOC have shown great potential when tested in such *in vitro* monolayer cultures, a significant decrease in their efficiency is commonly reported when progressed to *in vivo* settings. Tumour targeting with antibody-based therapies, such as ADCs, is a complex process in which not only antibody binding and internalisation efficiency, but also the penetration and diffusion ability of antibodies within the tumour 3D structure play a role in therapeutic efficiency (*16*). In addition, within the native tumour microenvironment (TME), tumour cells co-exist with stromal components such as natural killer cells and cancer-associated fibroblasts (CAFs), forming tight cell-cell and cell-matrix interactions (*17*). Consequently, with a view to selecting ADC candidates with higher chances of clinical success, better pre-clinical models that incorporate the 3D structure of the TME are needed.

In recent years, significant efforts have been put into the development of three-dimensional (3D) cultures, which better mimic the structure and behaviour of the native tumour (*15*) (*18*). Spheroids are emerging 3D systems, which consist of cellular aggregates made up of tumour cells alone or in the presence of other stromal components (*19*) (*20*) (*21*). Due to their compact nature, spheroids successfully recapitulate features of the tumour, such as its tight cell-cell interactions (*22*). Consequently, their use as models for the screening of novel drugs, such as ADCs, is of increasing interest. Advances in the 3D culture field have led to the optimisation of a range of high-throughput techniques which provide easy and reproducible protocols to grow spheroids in the lab (*23*). For instance, the liquid overlay technique is a method in which cells are seeded in culture plates pre-coated with compounds such as agarose, known as ultra-low attachment (ULA) plates, which prevent cell-plate interactions. Instead, they force cells to interact with each other to form spheroid aggregates (*14*).

In this study, we optimised ovarian cancer spheroid models which replicate the tight cell-cell features of the TME for the testing of ADCs. We demonstrated that epithelial adenocarcinoma cells from the human epidermal growth factor receptor 2 (HER2)-overexpressing cell line SKOV3 can form regular and reproducible spheroids when plated in ULA plates. Furthermore, after the optimisation of a range of imaging and drug treatment assays, we demonstrated that spheroid models are more suitable to study the potency and penetration dynamics of ADCs than monolayer cultures.

## MATERIALS AND METHODS

### Cell lines

Epithelial ovarian cancer cell lines SKOV3, OVSAHO, PEO1, PEA1, OVCAR4 and OVCAR3 were obtained from American Type Culture Collection (ATCC). Epithelial ovarian cancer Kuramochi cells were obtained from JCRB cell bank (Osaka, Japan). Cells were maintained in HEPES-containing RPMI 1640 medium (Sigma) supplemented with 10% Foetal Calf Serum (Eurobio) and 2 mM L-glutamine (LifeTechnologies). All cells were incubated at 37 °C in a humidified incubator at 5% CO2 atmosphere and passaged every 3-5 days when the confluence reached 70–80%. For passaging, the cells were washed once with phosphate buffered saline (PBS), incubated with Trypsin 0.25% -EDTA 1 mM (Sigma-Aldrich) for 5 min at 37 °C and replated at a 20-30% confluency.

### Spheroid cultures

Spheroid cultures were generated by seeding a defined number of cells onto Nunclon Costar ultra-low attachment (ULA) round bottom 96 well plates (Corning) in 100 µL of cell culture medium. After seeding, the ULA plates were centrifuged at 100 x g for 3 min, to allow for cell aggregation, and cultured at 37 °C and 5% CO2 atmosphere (Fig 1A). This procedure was used throughout the paper to grow spheroids regardless of the cell line type and the initial number of seeded cells.

**Figure 1.**
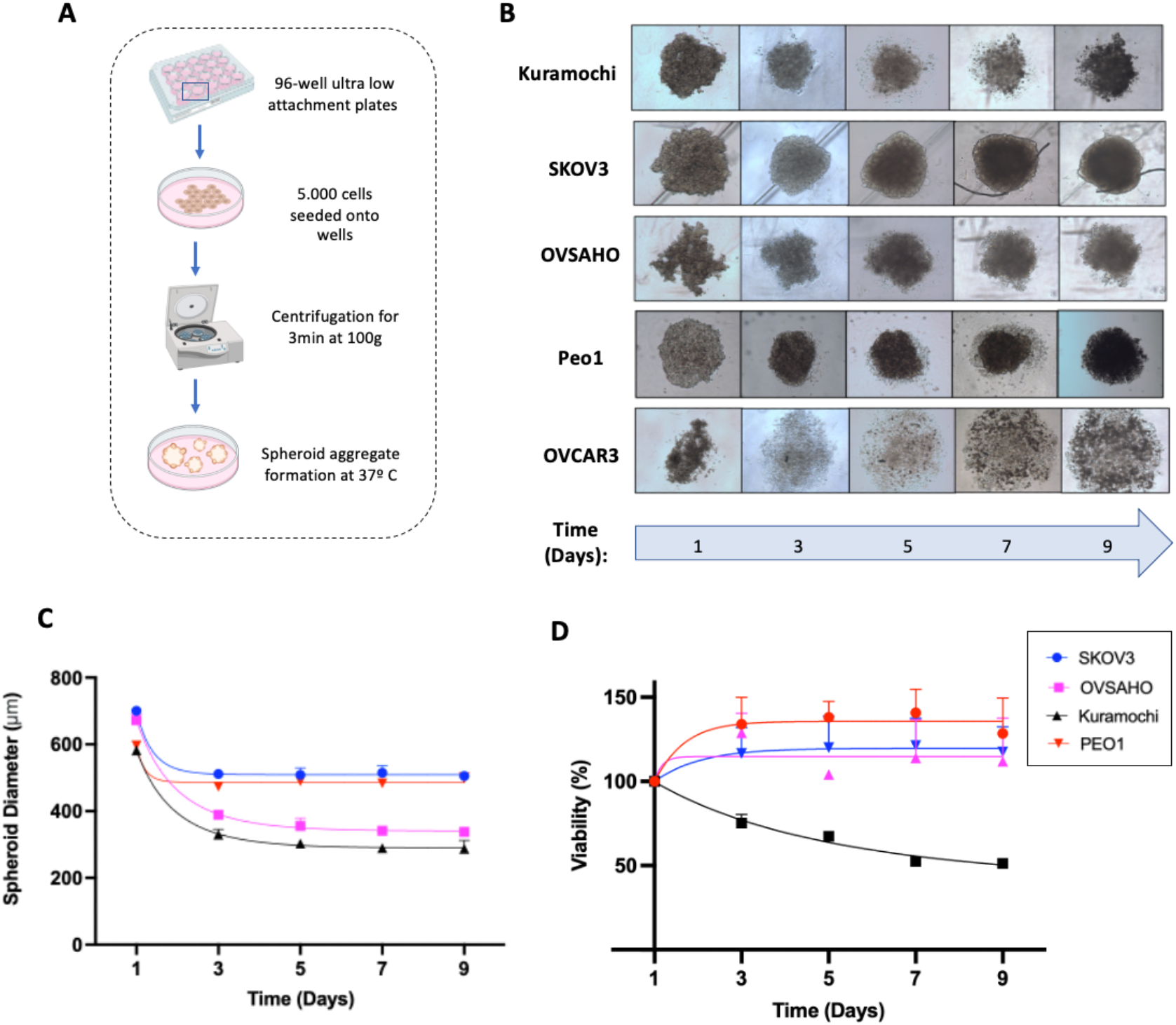
Optimisation of ovarian cancer spheroid growth. **A** Optimised protocol for the growth of spheroids in Ultra-low attachment plates. **B** Brightfield microscopy images of SKOV3, Kuramochi, OVSAHO, Peo1 and OVCAR3 spheroids of 5000 cells over 9 days. Day 0 is defined as the day of cell seeding. **C** Changes in the diameters of OVSAHO, Peo1 and SKOV3 spheroids over a course of 9 days. The diameter decrease of each spheroid is fit to the Gompertz equation. Data points are the mean volumes ± SD of four spheroids. **D** Growth viability of SKOV3, Kuramochi, OVSAHO and PEO1 spheroids tracked over 10 days. Data is expressed as the percentage viability relative to Day 1. Data points are the mean values ± SD of technical triplicates.

### Brightfield microscope imaging

OVSAHO, Kuramochi, PEA1, PEO1, OVCAR3 and OVCAR4 spheroids were generated following the cultivation strategy described above at densities of 5000 cells per well. SKOV3 spheroids were generated at densities between 2500 and 20000 cells per well. Over an incubation period of nine days, spheroids were imaged and captured using an inverted Nikon Diaphot-TMD light microscope with 4x/0.1 magnification lens, connected to a digital camera (micropublisher 5.0 RTV). To calculate the diameters of the captured spheroids, images were analysed using the ImageJ software.

HGSO cancer patient sections stained with the proliferation marker Ki-67 were obtained from our generous collaborators at Imperial College London. Images of the sections were captured using a light microscope with 10×/0.3 objective lens. The ImageJ software was used to calculate the diameter of the tumour islands.

### alamarBlue™ Cell Viability assay

Cellular viability of the ovarian cancer spheroids was studied over a nine-day period through the alamarBlue™ Cell Viability assay (Thermofisher). The Cell Titer Blue is an assay in which Resazurin is reduced to a highly fluorescent molecule, known as resorufin, in viable cells. To optimise the alamarBlue assay for its use on spheroids, 10 µL of alamarBlue® reagent was added directly to spheroids in 100 uL of culture and incubated in the dark at 37°C for 1, 2, 4 and 24 hours. As a negative control, 10 µL of alamarBlue reagent were added to wells with 100 µL of RPMI media alone. Fluorescence levels were measured using a spectrophotometer plate (Infinite® M1000 Pro, Tecan Group, Switzerland) at 560ex/ 590em and validated by visual comparison to the spheroid morphology. For data processing, the negative control fluorescence values were subtracted from the fluorescence values of experimental wells. Percentage viabilities across the culturing days were calculated relative to Day 1.

### Drug treatments with cytotoxic drugs and Antibody Drug Conjugates

Cytotoxic drugs, DM1 and MMAE, were obtained from Medchemexpress, and the primary antibody, trastuzumab was a generous gift from our collaborator at Oxford University. The antibody-drug conjugates, T-vcMMAE and T-DM1 were generated in Prof Eric Abouagye’s lab by Dr Ruisi Fu through chemical conjugation of trastuzumab with the MMAE and DM1 cytotoxic payloads, respectively.

SKOV3 spheroids of 5000 cells were generated in 96-well ULA plates following the cultivation strategy described above and incubated at 37°C. To generate monolayer cultures, SKOV3 cells were plated at a density of 500 cells/well in a flat bottom 96-well plate (Corning) in 100 µl of full RPMI media supplemented with 10% Foetal Calf Serum and 2 mM L-glutamine. On either the fifth or second day of spheroid and monolayer culture incubation, 100 µL of fresh RPMI medium containing 10% of penicillin in addition to a range of 4-fold increasing concentrations of DM1 (0 nM-400 nM), T-DM1 (0 nM-16 nM), MMAE (0 nM-4 nM) and Tvc-MMAE (0 nM-16 nM), were added to the cultures. The cultures were then incubated at 37°C and 5% CO2. Spheroids often have an initial growth phase of 3-5 days in which they progress from loose cell aggregates into compact spheroids that mimic the TME. When treated at the pre-mature stages of spheroid formation on day 2, the recorded viability data was shown to be highly inconsistent (data not shown). On the contrary, when treated on day 5 of spheroid culture, we obtained consistent data (Fig 3). For the purposes of this study, we therefore analysed the day 5 data. On the seventh day of drug exposure, the viability of these cultures was assessed and compared to untreated control.

### Viability assays for Drug-induced cytotoxicity analysis

Few viability assays have been optimised to accurately quantify drug-induced cytotoxicity in spheroids (*14*). Hence, the CellTiter-Glo 3D Cell Viability Assay (Promega) and the alamarBlue viability assays were assessed to measure drug-induced cytotoxicity in spheroids. The CellTiter-Glo 3D Cell Viability Assay is an assay in which cellular viability is measured through the recording of the levels of ATP. The alamarBlue assay was performed following the protocol described above. To perform the 3DGlo assay, a volume of 50 µL of the 3DGlo solution was added to the cell culture medium in each experimental well and in wells containing 200 µL of RMPI media. The media was pipetted up and down several times to ensure the rupturing of the spheroid aggregates. After a 10min incubation in the dark, 50 µL of the culture solutions in each well were transferred to opaque-walled 96-well plates and luminescence was recorded at a 1250 gain. The relative cell viability was calculated as a percentage of the viability of treated to untreated samples.

We determined that the 3DGlo assay gives viability data which is consistent and well-correlated with the visual reduction dynamics of the spheroid volumes upon treatment (data not shown). Consequently, the 3DGlo protocol was used throughout the study to investigate drug efficacy in spheroids.

### Flow Cytometry

SKOV3 spheroids of 5000 cells were generated in 96-well ULA plates as described above and cultured at 37°C. On their fifth day of culture, SKOV3 spheroids were treated with Trastuzumab (0.01 mg/mL) for 30 minutes, 4 hours, 24 hours and 72 hours. After Trastuzumab treatment, 12 spheroids were collected per condition, pooled together and washed in PBS. Single cell suspensions of each condition were obtained by incubating the collected spheroids in Trypsin 0.25% - EDTA 1 mM solution at 37 °C in 5% CO2 for 5 minutes. SKOV3 monolayer cultures were generated by seeding 10^6^ SKOV3 cells on 6 well-plates. As with spheroids, after 5 days of incubation, the monolayer cultures were treated with 0.01 mg/mL Trastuzumab for 30 minutes, 4 hours, 24 hours and 72 hours, washed on PBS and dissociated from the wells using Trypsin 0.25% - EDTA 1 mM solution. To perform an isotype control, SKOV3 spheroids and monocultures were also treated with 0.01 mg/mL of the non-specific binding antibody, Avastin Bevacizumab (Roche). The prepared cell suspensions were then washed in PBS twice to remove any excess antibody. Subsequently, cell suspensions were incubated for 30 min in ice with the fluorescently labelled secondary antibody, Alexa Fluor™ 488 goat anti-human IgG (Invitrogen), and with 4’,6-diamidino-2-phenylindole (DAPI), a staining for dead cells (Sigma). Following two additional PBS washing steps, the cell suspensions were analysed through flow cytometry using a FACSCalibur instrument (Becton Dickinson Franklin Lakes, NJ, USA). FITC fluorescence was excited using 488 nm argon laser, and DAPI fluorescence was excited using 640 nm laser. All results were analysed with FlowJo software (Tree Star).

To measure HER2 expression in OVSAHO and SKOV3 monolayer cultures, the above protocol was repeated. However, the cells were treated with 0.01 mg/mL Trastuzumab for 30 minutes in ice after their single cell suspensions were obtained. After two PBS washes, single cell suspensions were labelled with the non-specific binding antibody as above.

### Immunofluorescent microscopy and confocal imaging

To introduce a GFP tag in SKOV3 cells, second-generation lentiviral plasmids were obtained from Sinobiological and transfected into SKOV3 cells according to the manufacturer’s protocol. The lentiviral overexpression strategy was then validated through flow cytometry. The primary antibody, Trastuzumab, was conjugated with an Alexa FlourTM 633 fluorescent tag according to the manufacturer’s protocol (Invitrogen).

GFP-labelled SKOV3 spheroids of 5000 cells were generated as above and cultured at 37°. After the fifth day of growth, the spheroids were incubated for defined intervals of 30 minutes, 4 hours, 24 hours and 72 hours with 0.01 mg/mL of Alexa633-trastuzumab antibodies. The spheroids were then washed with PBS twice and transferred in full RPMI media to individual confocal chambers (ThermoFisher). Spheroids were imaged using an Inverted Lecia SP5 confocal microscope and Leica LAS AF Lite software. Widefield and fluorescent images of the spheroids were captured using the 10x water objective, the 488 nm laser diode for GFP excitation and the 640 nm laser diode for Alexa633 excitation.

Previous research has shown that due to the compact nature of spheroids, at certain z-planes, confocal imaging cannot fully penetrate and capture spheroid structures accurately (*24*). Consequently, the GFP-tag expressed by SKOV3 cells, was used as a reference to determine the z-planes at which the spheroids can be fully captured. Confocal z-slices of spheroids were acquired every 75-μm for a total of 150-μm from the top of the spheroids. Confocal z-slice images were analysed using the ImageJ software to obtain the average fluorescent intensity across the diameter of the spheroids.

### Statistical analyses

Data are expressed as Mean ± SD. Microsoft Excel and GraphPad Prism 7 (GraphPad Software, La Jolla, USA) were used for data analysis. Growth curves were obtained by fitting the data to the Gompertz growth model. Dose-response curves, area under the curve (AUC) values and half-maximal inhibitory concentration (IC_50_) values were calculated for each drug treatment by fitting the data to non-linear regression models. Statistical significances were analysed by paired students’ t test when appropriate. A p-value < 0.05 was considered as statistically significant and indicated as: *p<0.05, **p<0.01, ***p<0.005, p****<0.001, ns= non-significant

## RESULTS

### Production and optimisation of epithelial ovarian adenocarcinoma spheroids

To generate human ovarian adenocarcinoma spheroids, SKOV3, OVSAHO, Kuramochi, PEA1, PEO1, OVCAR3 and OVCAR4 cell lines were cultured for 9 days in ULA plates (Fig. 1A). The ability of each cell line to form spheroids was assessed by tracking changes in the growth dynamics, as well as the structural morphology of the plated cells.

Although all cell lines managed to quickly come together to form cell conglomerates within the first 24h of culture, these cell conglomerates acquired structurally different conformations later. At day 3, the SKOV3, PEO1, OVSAHO and Kuramochi cell conglomerates aggregated into spheroid-like structures of round borders and morphologies (Fig. 1B). Conversely, the OVCAR4, OVCAR3 and PEA1 cell lines disintegrated into non-defined and irregular cellular masses which do not feature spheroids (Fig. 1B). Consequently, the SKOV3, Kuramochi, OVSAHO and PEO1 cell lines were progressed on to further spheroid characterisation studies, whilst OVCAR4, PEA1 and OVCAR3 were disregarded.

To characterise the aggregation dynamics of the SKOV3, Kuramochi, PEO1 and OVSAHO spheroids, changes in their size were monitored over the period of incubation. The diameter of all four spheroids decreased throughout the first days of culture, until day 3, when spheroids stabilised to a constant size. The diameters of the PEO1, Kuramochi, SKOV3 and OVSAHO spheroids on day 7 of incubation were 438 mm, 289.5 mm, 514.5 mm and 341.5 mm, respectively.

Despite the observed similarities in the aggregation dynamics of Kuramochi, OVSAHO, PEO1 and SKOV3 spheroids, differences in their morphology and viability were evident throughout culturing (Fig. 1B and 1D). The viability assays revealed that the cellular viability of Kuramochi spheroids decreased progressively, whilst SKOV3, OVSAHO and PEO1 spheroids increased their cellular viability until day 9.

Additionally, we determined that the OVSAHO, Kuramochi and PEO1 spheroids do not have the tight compact morphology characteristic of spheroid models, due to their irregular borders and surrounding isolated cells (Fig. 1B). As SKOV3 spheroid was the only one that both increased its cellular viability and maintained a compact morphology, it was selected for further characterisation.

### Growth characterisation of SKOV3 spheroids

To study the clinical relevance of SKOV3 spheroid size, a range of cell densities were tested and tracked over a period of nine days. When below 2,500 cells per well, inconsistent spheroid growth was found. However, SKOV3 cells at cell densities between 2,500 and 20,000 cells per well were able to successfully form spheroids with similar morphologies (Fig. 2B). Whilst the volume of the lower density spheroids remained constant throughout 9 days, the volume of higher seeding spheroids decreased progressively until day 5 (Fig. 2A). On day 7, the spheroids with 2,500 cells per well had the smallest diameter of 483 µm. The spheroids with 20,000 cells per well had the largest diameter of 759.5 µm (Fig. 2A). Consequently, the sizes of SKOV3 spheroids were determined to be linearly dependent on their starting cell seeding number (Fig. 2B). These results indicate that SKOV3 cells can form spheroids at a range of cell densities, which can be exploited to obtain spheroid sizes at the user’s convenience.

**Fig. 2.**
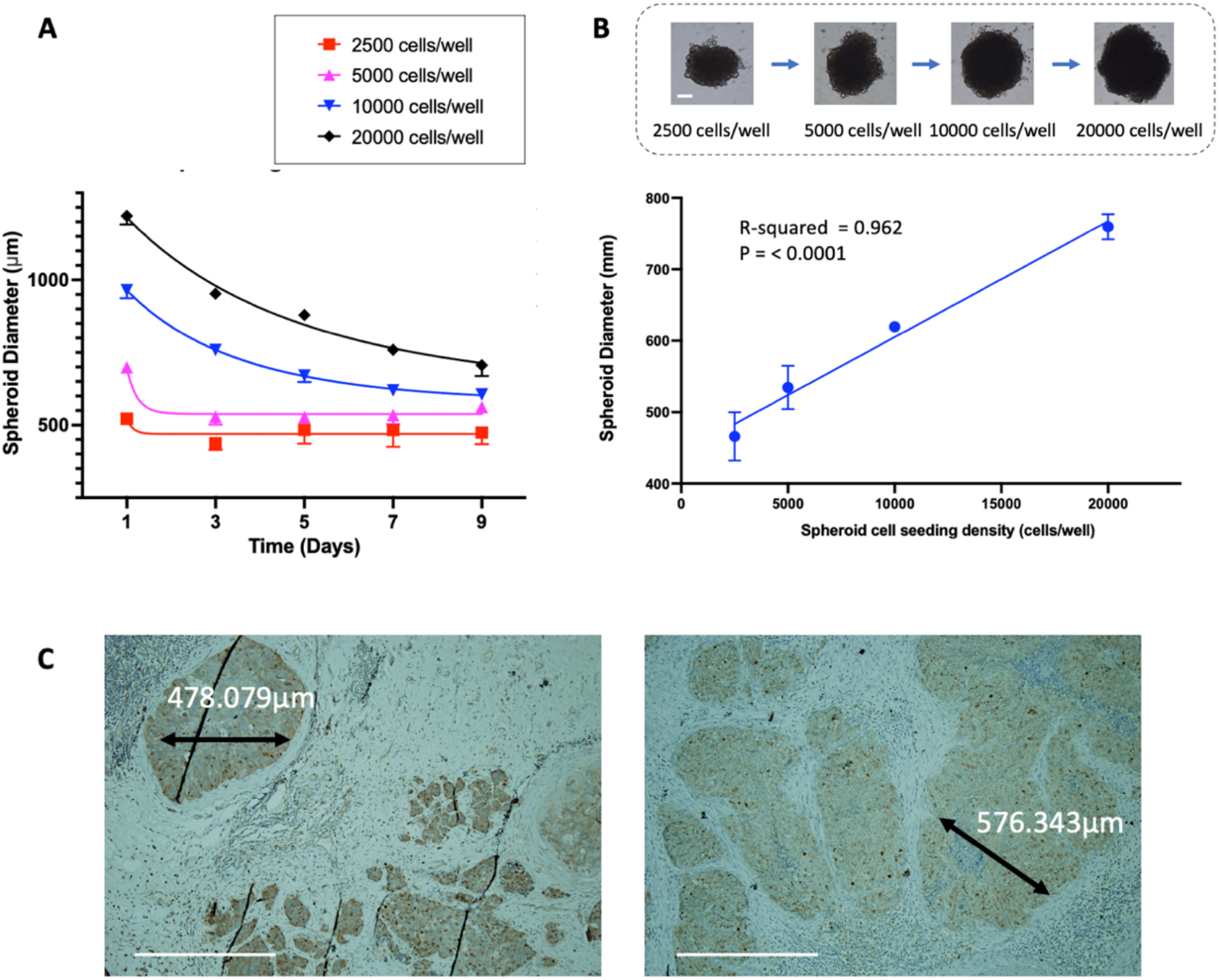
SKOV3 spheroid growth kinetics and morphology. **A** Changes in the diameter of SKOV3 spheroids of different cell seeding densities (2500, 5000, 10000 and 20000 cells/well) over 9 days. The diameter decrease data for each spheroid size is fit using the Gompertz equation. Data are presented as means ± SD of technical triplicates. **B** Linear relationship between the cell plating densities of SKOV3 spheroids and their size on day 7 of culture. Simple linear regression is fit through the data. Data are presented as means ± SD of 3 spheroids per experiment. Images are from the same spheroids. **C** Brightfield microscopy images of histological sections from HGSO cancer patients stained with Ki-67 (brown). Representative sizes of tumour islands are indicated by the black arrows. Scale bar (white): 500 µm. Only two histological sections are shown for simplification (n=5).

With a view to determining the clinical relevance of SKOV3 spheroids, their diameter was compared to the sizes of the tumour islands within the patients’ tumour tissues. The diameters of the tumour islands ranged from 400 µm to 800 µm (Fig. 2C). As SKOV3 cells plated at 5000 cells per well formed compact and reproducible spheroids of similar diameters to those found in the tumour islands, they were selected for downstream ADC sensitivity characterisation.

### Cytotoxicity of antibody-drug conjugates in spheroids vs monolayer cultures

As confirmed by flow cytometry (Fig. 3A), SKOV3 cells express high levels of HER2, a surface protein that is commonly overexpressed in breast cancers and a subset of ovarian cancer patients. Hence, to investigate the impact of monolayer and spheroid cultures on ADC sensitivity, cell viability was assessed following treatment with two HER2 targeting ADCs: T-DM1 and T-vcMMAE.

**Figure 3.**
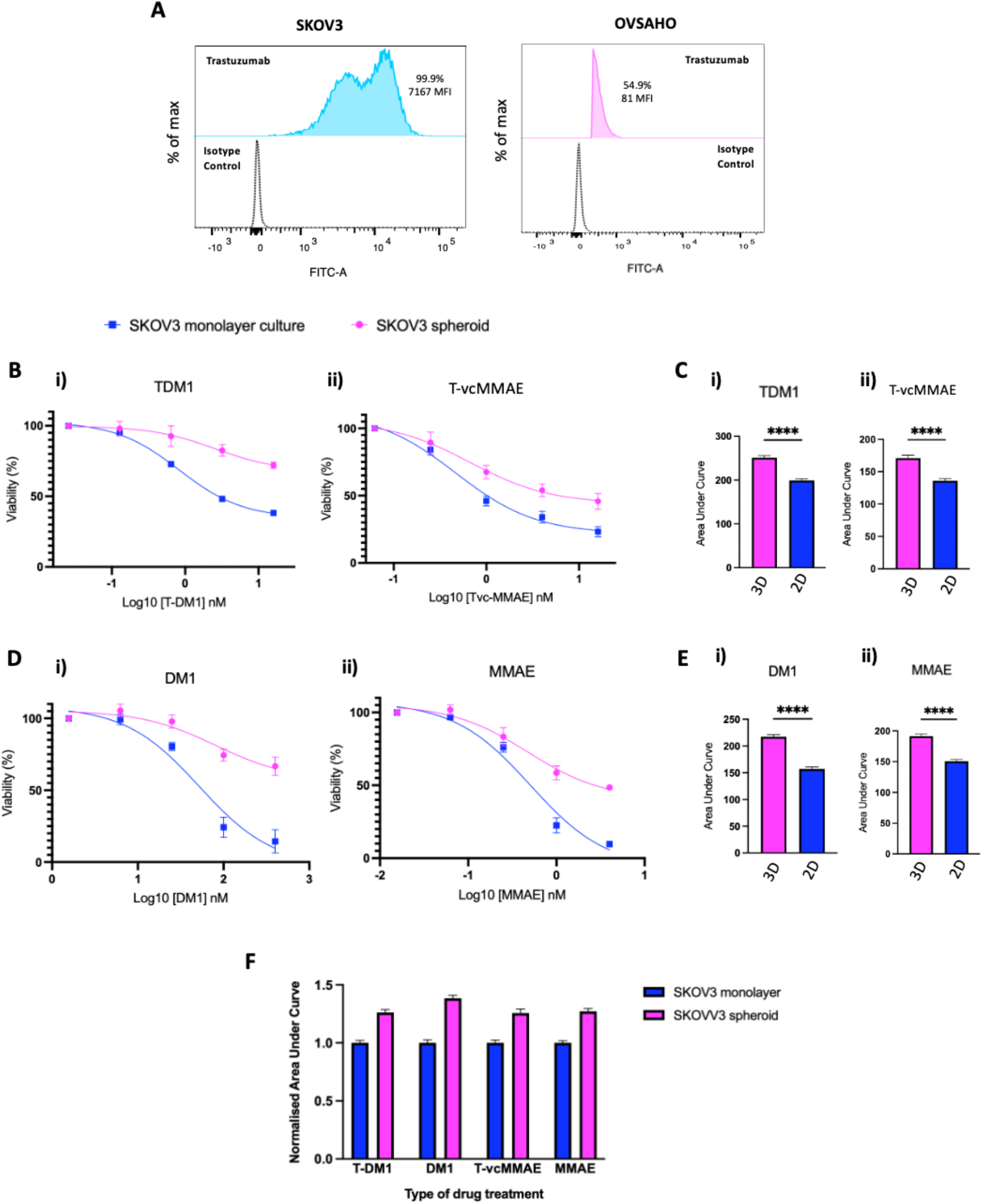
Cytotoxicity effects of ADCs and cytotoxic drugs in 3D spheroid cultures vs 2D cultures. **A** Analysis of HER2 expression in SKOV3 cells. The distribution of SKOV-3 and OVSAHO cells (negative control) are presented according to their FITC intensity after incubation with HER2-specific antibody trastuzumab or isotypic control antibody. The presented data is representative of 2 independent experiments showing similar results. **B** Concentration-dependent viability changes of SKOV3 spheroids and monolayer cultures after 7-days of treatment with 4-fold increasing concentrations of i) T-DM1 and ii) T-MMAE. Data is expressed as the percentage viability relative to untreated control and fit to a non-linear regression model. Data points are presented as means ± SD of 4 individual experiments, performed in technical triplicate. **C** Area under the curve values of the i) T-DM1 and ii) T-MMAE dose-dependent curves in SKOV3 monolayer and spheroid cultures. Statistical significance is calculated using a two-tailored Student’s t-test (****P<0.001). **D** Concentration-dependent viability changes of SKOV3 spheroids and monolayer cultures after 7-days of treatment with 4-fold increasing concentrations of i) DM1 and ii) MMAE. Data is expressed as the percentage viability relative to untreated control and fit to a non-linear regression model. Data points are presented as means ± SD of 4 individual experiments, performed in technical triplicate. **E** Area under the curve values of the i) DM1 and ii) MMAE dose-dependent curves in SKOV3 monolayer and spheroid cultures. Statistical significance is calculated using a two-tailored Student’s t-test. ****P<0.001. **F** AUC values of ADCs and cytotoxic dose-dependent curves normalized against monolayer culture for each drug treatment.

Following 7 days of T-DM1 and T-vcMMAE treatment, dose-dependent curves of the monolayer and spheroid cultures were obtained (Fig. 3B). Although the cell viability of both the monolayer and spheroid decreased progressively in response to increased ADC concentrations, spheroid culture yielded significantly lower levels of sensitivity. In monolayer cultures, T-vcMMAE and T-DM1 yielded area under the curve (AUC) values of 136 and 199, respectively, and IC_50_ values of 1.02 nM and 3.2 nM, respectively. In spheroid cultures, T-vcMMAE and T-DM1 yielded AUC values of 170.9 and 251.2, respectively. As the SKOV3 spheroids did not reach ≥50% reductions in cell viability, their IC_50_s could not be estimated (Fig. 3C). The differences between the AUC values of T-vcMMAE and T-DM1 in 2D and 3D cultures were statistically significant with p-value < 0.001 (Fig 3C). These results thus suggest that SKOV3 spheroids are less responsive to treatment with ADCs than SKOV3 in monolayer culture cells.

ADCs contain an antibody of a large molecular weight (150 kDa) as well as cytotoxic payloads of lower molecular weights (<1 kDa). Previous studies have hypothesised that antibodies have difficulties in penetrating tumour-like spheroids (*25*). Hence, to determine whether the size of T-DM1 and T-vcMMAE contributes to a reduced sensitivity in spheroids, the potency of their cytotoxic payloads, DM1 and MMAE, were assessed in spheroid and monolayer cultures (Fig. 3D). Surprisingly, spheroids also showed reduced sensitivities to both cytotoxic payloads. DM1 treatment yielded AUC values of 217.4 in spheroids and 157 in monolayer cultures. Furthermore, MMAE yielded AUC values of 191.6 in spheroids and 150.7 in monolayer cultures. The differences between the AUC values of DM1 and MMAE in 2D and 3D cultures were statistically significant with p-value ****< 0.001 (Fig. 3E). Furthermore, while all treated monolayer cultures reached their IC50 values, the SKOV3 spheroids did not.

Additionally, compared to T-DM1, MMAE and T-MMAE, DM1 showed the highest levels of sensitivity differences between spheroid and monolayer cultures (Fig. 3F). These results indicate that the size of the ADC molecules may not be the main reason for their reduced sensitivity in spheroids.

### Dynamics of ADC Penetration into Spheroids

Since both ADCs and cytotoxic payloads exhibited similar degrees of resistance in spheroids compared to monolayer cultures, we aimed to study the penetration dynamics of large molecules into spheroids. In particular, we assessed the penetration dynamics of trastuzumab, the antibody component of T-DM1 and T-MMAE.

SKOV3 spheroids and monolayer cultures were incubated with trastuzumab between 30 minutes and 72 hours, and the cell populations that were accessed and bound by the antibody were analysed through flow cytometry. In monolayer cultures, cell populations started to show high FITC intensities at 30 minutes, which remained throughout 72 hours of treatment (Fig. 4A). These findings suggest that trastuzumab can access and bind most SKOV3 cells in monolayer as early as 30 minutes of incubation. Notably, FITC intensity was reduced slightly at 72 hours compared to 24 hours, possibly due to internalisation of trastuzumab at 72 hours (Fig. 4A).

**Figure 4.**
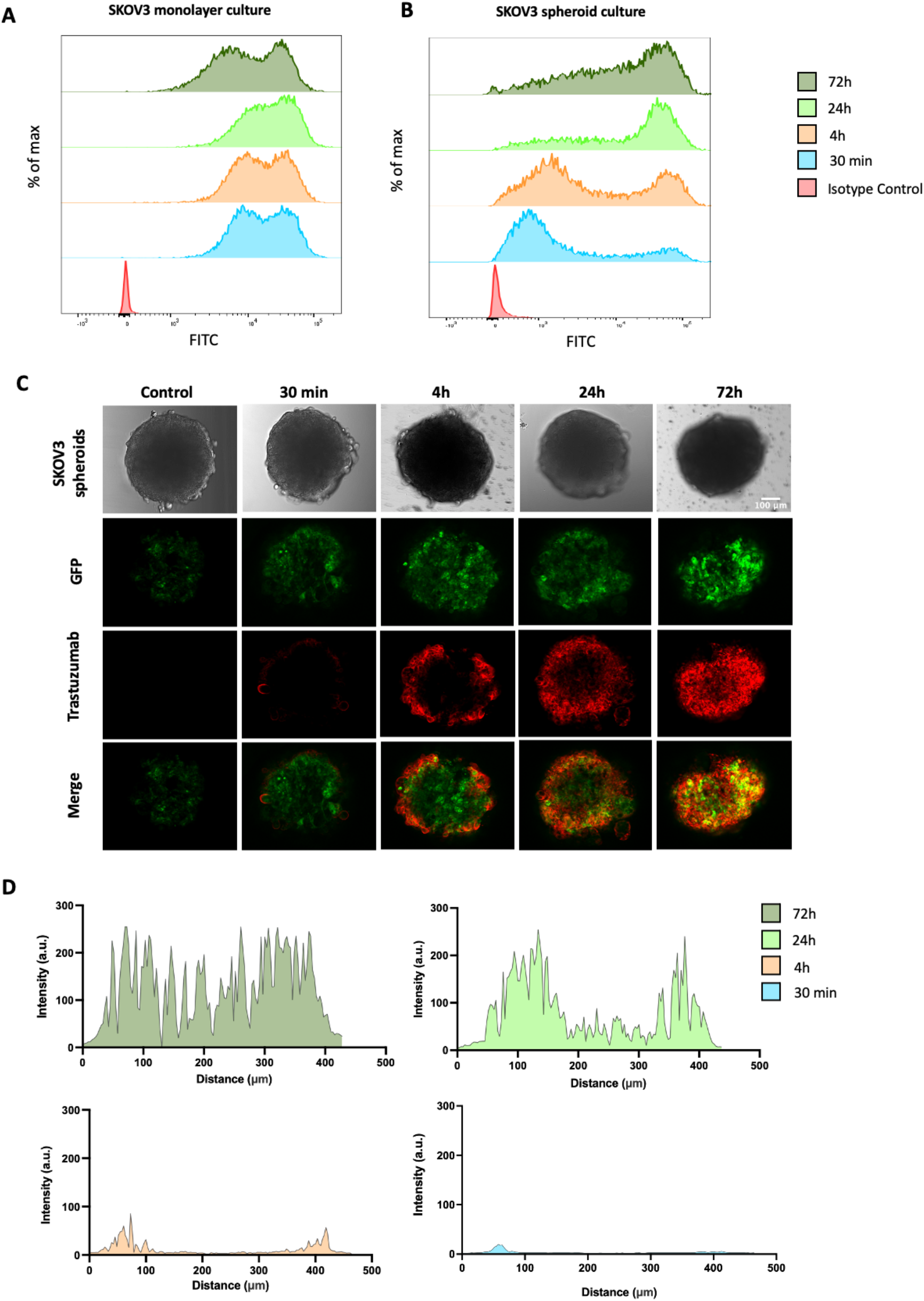
Penetration dynamics of Trastuzumab in SKOV-3 spheroids. Distribution of SKOV-3 (**A**) monolayer and (**B**) spheroid cells according to their FITC intensity after trastuzumab treatment for up to 72 hours. The histograms in red represent the isotype control. The presented data is representative of 3 independent experiments showing similar results. **C** Immunofluorescence microscopy images of GFP-labelled SKOV3 spheroids treated with and without Alexa633-labelled Trastuzumab for up to 72 hours. The fluorescence signal of GFP and Alexa633 is presented in green and red, respectively. Images are taken at the same z-plane (75 μm from the top of the spheroids). Images are representative of 2 independent experiments with technical duplicates each, showing similar results. **D** Fluorescence intensity across the diameter of the imaged GFP-spheroids treated with Alexa633-Trastuzumab for 30 minutes, 4 hours, 24 hours and 72 hours.

In contrast to the monolayer cultures, cell populations in spheroids showed low FITC intensity at 30 minutes and 4 hours, indicating low levels of trastuzumab access and binding (Fig 4B). However, after 24 hours of incubation, the cell populations shifted to higher FITC intensities, similar to those recorded in monolayer cultures. This suggests that trastuzumab takes significantly longer to penetrate spheroids compared to monolayer cultures, which may partly contribute to the ADC resistance in spheroids.

To validate the flow cytometry findings, the penetration efficiency of trastuzumab across spheroids was also assessed through immunofluorescence microscopy. GFP-tagged SKOV3 spheroids were treated with AlexaFluor633-conjugated trastuzumab antibodies for a range of time points (30 minutes, 4 hours, 24 hours and 72 hours) and visualised using confocal lasers. Consistent with the flow cytometry experiments, trastuzumab was shown to gradually penetrate through spheroids over time (Fig. 4C). At 30 minutes and 4 hours, the fluorescence signal was only detected on the outer layer of the spheroids. However, this signal started to accumulate inside the spheroids from 24 hours (Fig. 4D). In fact, most cells in spheroids seemed to have successfully bound to the antibody at 72 hours.

## DISCUSSION

It has been hypothesised that the *in vivo* potency of antibody-based therapies, such as antibody-drug conjugates (ADCs), is influenced by their ability to penetrate, diffuse and interact with the three-dimensional tumour microenvironment (TME) (*17*) (*26*). Currently, ADCs under pre-clinical development against EOC are tested in two-dimensional *in vitro* cultures which do not mimic the native tumour (*27*). In this study, we have established and optimised a 3D epithelial ovarian cancer spheroid model as well as a range of analysis and imaging assays for ADC development.

Since the establishment of the first spheroid model in 1970, there has been significant progress in the development of methodologies for the growth of 3D cultures (*14*). Ultra-low attachment plates are platforms which force cells to interact with each other and aggregate into spheroids (*15*) (*28*). In this study, we showed that, when plated in ULA plates, different ovarian cancer cell lines form spheroids at different efficiencies. Whereas the SKOV3, Kuramochi, OVSAHO and PEO1 cell lines successfully aggregated into spheroid-like structures, PEA1, OVAR4 and OVCAR3 did not. Such significant differences in the ability of cell lines to form spheroids suggest that they have distinct aggregation and compaction dynamics. According to transcriptomics analysis performed by other groups, cancer cell lines express adhesion molecules aberrantly (*29*) (*30*), which is likely to result in differences when forming spheroids. For future work, transcriptomics analyses could be performed to identify the genotypic features which promote the successful formation of spheroid models. This work could aid the process of automating the selection of cell lines for spheroid growth.

Of the four cell lines that aggregated into spheroid-likes structures, we observed that SKOV3 cells formed the most compact and regular structures. Consequently, the SKOV3 cell line was progressed onto further spheroid characterisation studies. We showed that within the initial days of culturing, SKOV3 cells formed tight interactions with each other which led to a decrease in their volume. Given that this decrease in size was accompanied by increased levels of cellular viability, we hypothesise that SKOV3 spheroids become more and more cellularly compact over time. Previous groups have shown that due to their compact nature, spheroids are made up of a range of layers that mimic features of the native TME: a proliferative external layer, a quiescent intermediate layer and an inner hypoxic core, characterised by nutrient and oxygen deficiency (*15*) (*31*). Because of the lack of vasculature within our established SKOV3 spheroids, their core is likely to reproduce the hypoxic environment of the native tumour. Furthermore, due to their high proliferation rates, we also hypothesise that they are likely to maintain the external proliferative layer. However, to validate these findings, further structural and gene expression studies are required.

In this study, we also showed that SKOV3 cells formed compact and reproducible spheroids of sizes that were linearly dependent on their cellular density (*32*). For our studies to be clinically relevant, the SKOV3 spheroid models that had similar sizes to tumour islands present within patient tumour tissues were selected for further studies.

The aforementioned results suggest that SKOV3 spheroids mimic the tight cell-cell interactions, as well as the compact structure of the TME more efficiently than monolayer cultures. To assess the role the compact nature of the TME plays in ADC efficiency, the SKOV3 spheroid model was progressed to drug treatment studies. As the SKOV3 cell line expresses high levels of HER2, a range of viability and imaging studies were optimised in SKOV3 spheroids using the HER-2 specific ADCs, Ado-trastuzumab emtansine (T-DM1) and trastuzumab-monomethyl auristatin E (T-vcMMAE) (*33*). T-DM1 is an ADC which has been clinically approved for the treatment of HER2-overexpressing breast cancer (*13*) (*34*). T-vcMMAE is another ADC that, although not yet approved, is also under clinical development for breast cancer treatment (*35*). T-DM1 and T-vcMMAE are composed of the HER2-specific antibody, trastuzumab, and the cytotoxic payloads, DM1 and MMAE, respectively.

The cytotoxic studies optimised and performed within this paper showed that SKOV3 spheroids exhibited higher resistance to T-DM1 and T-vcMMAE compared to monolayer cultures. Previous studies investigating ADC behaviour have hypothesised that large molecular weight molecules such as ADCs have low penetration efficiencies when crossing compact surfaces (*36*). Furthermore, several groups have also suggested that, when ADC-specific antigens are highly expressed by tumour cells, the antigens form a barrier that complicates antibody penetration (*22*). Consequently, we initially attributed the lower potency of ADCs in spheroids to these 3D-associated penetration limitations. Unexpectedly, we then observed that, similarly to ADCs, their cytotoxic payloads, DM1 and MMAE, also yield lower potencies in spheroids compared to monolayer cultures. These payloads are of much smaller size than ADCs and, furthermore, do not kill tumour cells by specifically binding to HER2 antigens. Consequently, these findings suggested that the large size of the ADC molecules may not be the main reason behind their reduced sensitivity.

After the optimisation of a range of imaging studies in spheroid cultures, we showed that ADCs are able to penetrate through spheroids within 24 hours of treatment. Other groups have shown that molecules of low molecular weight, such as DM1 and MMAE, penetrate compact surfaces at much faster rates than large molecules of high molecular weight (*32*) (*37*) (*38*). For instance, Zhang *et al*., recently showed that if the molecular weight of the molecular drug carrier, octyl chitosan (OC), is 1kDa, it is able to fully penetrate into spheroids after 8 hours of incubation (39). However, if the molecular weight of OC is of 10kDa, its penetration efficiency drops by over 50% after 8h of incubation. Given these observations and the fact that T-MMAE and T-DM1 have a molecular weight of 150 kDa, whilst their cytotoxic payloads have a molecular weight of 1kDa, we suggest that ADCs are likely to penetrate spheroids more slowly than their payloads. A limitation of these findings, however, is that we did not experimentally analyse the exact penetration dynamics of the cytotoxic payloads alone. To validate these findings, future studies could determine the penetration dynamics of small molecules relative to ADCs through the optimised spheroids.

As ADCs consist of an antibody backbone attached to a number of cytotoxic payloads, once ADCs have reached a tumour cell, they deliver several cytotoxic drugs. T-DM1 and T-vcMMAE have a drug to antibody ratio (DAR) of ∼8. Therefore, we hypothesise that even if both ADCs penetrate through spheroids more slowly than their payloads, they could have comparable capability to kill cancer cells in 3D structure due to their high drug-to-antibody ratios. Further experiments to test efficacy of ADCs with a range of DAR in sperhoids are required to test this hypothesis.

Another limitation of this study is that all cytotoxicity and imaging studies were performed on spheroids grown from SKOV3, a cell line that originates from the ascetic fluid of a patient with ovarian cancer (*40*). As this cell line is not derived from the “primary tumour”, extensive evidence has highlighted that it does not accurately represent the genotypic and phenotypic features of epithelial ovarian cancers (*40*). Consequently, future studies are required to replicate the experiments in cell lines derived from the primary tumour of ovarian cancers. Although a range of ovarian cancer cell lines were tested in this paper for spheroid growth, they failed to form compact spheroid structures. Other groups have determined that extracellular components such as methylcellulose or matrigel promote the formation of compact and round spheroids, which mimic more realistic tumour microenvironment (*41*) (*27*) (*42*). For future ADC characterisation studies, ovarian adenocarcinoma cell lines could be tested for spheroid growth in the presence of different extracellular components.

Extensive evidence has shown that there are two main mechanisms of ADC-targeted tumour killing: through the aforementioned release of the cytotoxic payloads within tumour cells, and through the binding and recruitment of NK cells to the native tumour(*10*). When recruited by ADCs, NK immune cells signal tumour cells to die through the activation of a mechanism known as antibody dependent cellular cytotoxicity (ADCC) (*12*). With a view to studying the role the ADCC pathway plays in ADC responses, future work could focus on establishing and optimising tumour-immune co-culture spheroid systems.

In this study, we have optimised a range of EOC imaging and analysis assays which could be exploited for the development of novel ADC candidates against EOC. We suggest that due to the role the compact tumour 3D structure plays in therapeutic resistance, spheroids are more suitable models to study the potency and penetration dynamics of ADCs than monolayer cultures.

## ABBREVIATIONS

EOC: epithelial ovarian cancer;
ADCs: Antibody drug conjugates;
TME: Tumour microenvironment;
ULA: ultra low attachment;
ADCC: antibody dependent cell-mediated cytotoxicity;
HER2: human epidermal growth factor receptor 2;
SD: Standard deviation;
FITC: Fluorescein isothiocyanate;
AUC: area under the curve;
Cancer: associated fibroblasts,CAFs;
NK cells: natural killer cells;
T-DM1: Ado-trastuzumab emtansine;
T-vcMMAE: Trastuzumab-monomethyl auristatin E

## ACKNOWLEDGMENT

The authors acknowledge the support from Air Theranotiscs Ltd for this work.

